# An aphid effector promotes barley susceptibility through suppression of defence gene expression

**DOI:** 10.1101/639476

**Authors:** Carmen Escudero-Martinez, Patricia A. Rodriguez, Pablo A. Santos, Jennifer Stephens, Jorunn I.B. Bos

## Abstract

Aphids secrete diverse repertoires of effectors into their hosts to promote the infestation process. While “omics”-approaches facilitated the identification and comparison of effector repertoires from a number of aphid species, the functional characterization of these proteins has been limited to dicot (model) plants. The bird cherry-oat aphid *Rhopalosiphum padi* is a pest of cereal crops, including barley. Here, we extended efforts to characterize aphid effectors with regards to their role in promoting susceptibility to the *R. padi-*barley interaction. We selected 3 *R. padi* effectors based on sequences similarity to previously characterized *M. persicae* effectors and assessed their subcellular localisation, expression, and role in promoting plant susceptibility. Expression of *R. padi* effectors RpC002 and Rp1 in transgenic barley lines enhanced plant susceptibility to *R. padi* but not *M. persicae*, for which barley is a poor host. Characterization of Rp1 transgenic barley lines revealed reduced gene expression of plant hormone signalling genes relevant to plant-aphid interactions, indicating this effector enhances susceptibility by suppressing plant defences in barley. Our data suggests that some aphid effectors specifically function when expressed in host species, and feature activities that benefit their corresponding aphid species.

## Introduction

Similar to plant pathogens, aphids form close associations with their hosts and secrete effector molecules to modulate host cell processes to their benefit. Over the past decade, a combination of genomics- and proteomics-based approaches allowed the identification of putative effectors from different aphid species, including economically important pests of both monocot and dicot crops (Nicholson *et al*., 2012) (Rao *et al*., 2013) (Cooper *et al*., 2011) (Atamian *et al*., 2013) (Vandermoten *et al*., 2014) (Thorpe *et al*., 2016) (Zhang *et al*., 2017) (Thorpe *et al*., 2018). Comparative analyses of aphid effector repertoires across species has revealed core and diverse sets, provided insight into effector diversity, and evidence for a shared transcriptional control mechanism driving their expression (Thorpe *et al*., 2016) (Boulain *et al*., 2018) (Thorpe *et al*., 2018). Moreover, functional characterization of aphid effectors increased our understanding of how these proteins may function to enhance plant susceptibility during infestation (as reviewed by (Yates and Michel, 2018) and (Nalam *et al*., 2019)), and pointed to host-specific effector activities (Pitino and Hogenhout, 2013) (Elzinga *et al*., 2014) (Rodriguez *et al*., 2017b).

The C002 salivary protein was first described as an effector in *Acyrthosiphon pisum*, and promotes host susceptibility to aphids (Mutti *et al*., 2008). However, while expression of MpC002 (*M. persicae* C002) in host species *Arabidopsis* and *N. benthamiana* enhances susceptibility to *M. persicae*, expression of ApC002 from *A. pisum* in these same plant species has no visible impact on the host interaction with *M. persicae* (Pitino and Hogenhout, 2013). The difference in effector activity was attributed to a motif sequence (NDQGEE) in the N-terminal region of MpC002, which is lacking in ApC002 (Pitino and Hogenhout, 2013). In addition, several effectors from the broad host range pest *M. persicae* have been implicated in promoting host susceptibility, including Mp1 and Mp58 (Pitino and Hogenhout, 2013) (Elzinga *et al*., 2014) (Rodriguez *et al*., 2017b). However, the underlying mechanisms by which these effectors impact susceptibility remain largely unknown. We previously described that Mp1 associates with the host trafficking protein VPS52 (Vacuolar Sorting Associated Protein 52) to promote plant susceptibility (Rodriguez *et al*., 2017a). Using different combinations of Mp1 and VPS52 variants from different plant and aphid species, respectively, we showed that the Mp1-VPS52 association is highly specific to the broad host range pest *M. persicae* and its hosts, and likely shaped by plant-aphid co-evolution. Critically, effector-host protein interactions correlate with effector virulence activities (Rodriguez *et al*., 2017a). The Mp1 and Mp58 effectors and their putative orthologs are genetically linked across the genomes of at least 5 different aphid species (Thorpe *et al*., 2018). Although a functional link between Mp1 and Mp58 remains to be elucidated, Mp58 was previously implicated in plant-aphid interactions. For example, Elzinga *et al*. 2014 observed a decrease in *M. persicae* performance when Mp58 was ectopically expressed in *Nicotiana tabacum* or transgenic Arabidopsis lines. In contrast, the Mp58-like effector from *M. euphorbiae* (also called Me10) enhances tomato and *N. benthamiana* susceptibility to *M. euphorbiae* and *M. persicae* (Atamian *et al*., 2013). Me10 was recently reported to interact with tomato 14-3-3 isoform 7 (TFT7), which contributes to defence against aphids (Chaudhary *et al*., 2019).

*Rhopalosiphum padi* is an aphid species with a narrow host range, which includes grass species, such as barley, oats and wheat (Blackman and Eastop, 2000). This aphid is an important pest of cereal crops that causes feeding damage and transmits some of the most destructive viral diseases of cereals, such as *Barley Yellow Dwarf Virus* (BYDV). Whilst *R. padi* is highly specialized on cereals, other species, like *M. persicae*, feature an exceptionally broad host range that includes more than 4000 different plant species (Blackman and Eastop, 2000), including the model plants Arabidopsis or *N. benthamiana*. Despite its broad host range, *M. persicae* is not a pest of barley and performs poorly on this plant species (Escudero-Martinez *et al*., 2017). Recently, *M. persicae* and *R. padi* effector repertoires were identified and compared, allowing the extension of effector characterization studies to cereal pests (Thorpe *et al*., 2016; Thorpe *et al*., 2018). Functional characterization of aphid effectors across different plant species, including cereals is important to gain insight into sequence variation among effector repertoires impacts host susceptibility.

Here, we characterized 3 *R. padi* effectors with regards to their subcellular localization, gene expression, and contribution to susceptibility in host barley and non-host *N. benthamiana* plants. We found that expression of the *R. padi* effectors Rp1 and RpC002 in transgenic barley lines enhances plant susceptibility to *R. padi* (host interaction) but not to *M. persicae* (poor-host interaction), highlighting the importance of these effectors for barley colonization in an aphid species-specific manner. Further characterization of Rp1 transgenic barley lines revealed reduced expression of several markers of plant hormone signalling pathways relevant to plant-aphid interactions, suggesting this effector may enhance susceptibility by suppressing plant defences.

## Material & methods

### Aphid cultures

Aphids used for the experiments were raised inside cages under controlled conditions in growth chambers (18°C, 16h light). *R. padi* was raised on *Hordeum vulgare* L. cv. Optic and *M. persicae* (genotype O) was reared on *Brassica napus*. The aphid species used were kindly provided by Alison Karley (JHI, UK) and Gaynor Malloch (JHI, UK).

### Identification of putative effector orthologs and plasmid construction

Effector annotation and identification of orthologs was performed as described by Thorpe *et al*. 2016. Similarity searches were performed by reciprocal best BLAST hit analysis between *R. padi* and *M. persicae* transcriptomes with the minimum thresholds of 70 % identity and 50 % query coverage. Pair-wise sequence analysis was performed in Jalview 2.10.4 (Waterhouse *et al*., 2009) with T-coffee and default parameters. Signal peptide sequences were predicted with SignalP 4.1 (Petersen *et al*., 2011). Coding sequences were amplified from *R. padi* and *M. persicae* cDNAs, without the region encoding for the signal peptide, and verified by sequencing (Primers in Supplementary Table S1). The resulting amplicons were cloned by Gateway technology into pDONR201, pDONR207 or pENTR_D-TOPO (Gateway®, Invitrogen). Sequence verified inserts were cloned into different destination vectors by LR reaction. Destination vectors pB7WGF2 (35S promoter, N-terminal GFP) and pB7WG2 (35S promoter, no tag) (Karimi *et al*., 2002) were used for transient overexpression in *N. benthamiana*, and pBRACT214m (maize ubiquitin promoter, no tag), kindly provided by Abdellah Barakate (JHI) (Colas *et al*., 2019), was used for generating transgenic barley lines.

### Effector gene expression in aphids exposed to host-, non-/poor-host plants and artificial diet

The experimental set-up for determining aphid effector gene expression in aphids exposed to the different feeding environments is explained in detail in Thorpe et al., (Thorpe *et al*., 2018). Briefly, aphids were exposed to an artificial diet, host, poor/nonhost plant for 3h and 24h and collected for RNA samples preparation and their transcriptome was sequenced by RNAseq. More specifically, *R. padi* was exposed to barley (host) and Arabidopsis (non-host), and *M. persicae* was exposed to Arabidopsis (host) and barley (poor-host). Both aphids were also exposed to artificial diet for 3h and 24h. A total of five independent replicates were used for this experiment and differential expression (DE) analyses was performed as described (Thorpe *et al*., 2018). For each selected effector (Rp1, RpC002, Rp58, Mp1, MpC002, and Mp58), we performed BLAST searches against the RNAseq datasets described in Thorpe *et al*. (2018) to identify their corresponding gene models.

Transcripts were normalized by the fragments per kilo-base of exon per million reads mapped (TMM-FPKM) method, which normalized the gene counts to the gene length and the library size (Conesa *et al*., 2016).

### Effector localization

Effectors were cloned into pB7WGF2 and the constructs were transformed into *Agrobacterium tumefaciens* strain GV3101. *Agrobacterium* cells were harvested by centrifugation (8min, 6000rpm) and resuspended in infiltration buffer (acetosyringone 125 µM and MgCl_2_ 10mM) to an optical density of OD_600_= 0.1. *Agrobacterium* carrying the GFP-effector constructs were then infiltrated in *N. benthamiana* leaves. RpC002 and MpC002 were co-expressed with a plasma membrane molecular marker (Nelson *et al*., 2007) and with the p19 silencing suppressor (OD_600_ = 0.1) for improving expression and thereby detection under the confocal microscope. All other effector pairs were infiltrated without p19. Three days after infiltration, plants were analysed under the confocal microscope (Zeiss LSM 510 (Jena, Germany) using a Zeiss x20/x40 lens. GFP-fusion proteins and free GFP were imaged with the GFP and chlorophyll filter (488 nm excitation), and co-localization with the membrane marker was analysed with GFP-RFP filter (488-561nm excitation). The experiment was repeated four times, and the resulting images were processed using ImageJ (Schneider *et al*., 2012).

### Western blotting to detect GFP-fusion proteins

Effectors were cloned into the pB7WGF2 vector and constructs were transformed into *Agrobacterium tumefaciens* strain GV3101. *Agrobacterium* cells were treated as above and infiltrated in *N. benthamiana* leaves to an optical density of OD_600_ = 0.3. After four days, samples were harvested, and proteins were extracted with GTEN buffer (10% Glycerol, 25mM Tris pH 7.5, 1mM EDTA, 150mM NaCl, 0.1% NP-40, 10mM DTT and 1x protease inhibitor cocktail, Sigma). Western blots were incubated overnight with GFP-antibody (Santa Cruz Biotechnology Inc, USA), for 1 hour with anti-rabbit-HRP (Santa Cruz Biotechnology Inc, USA).

### Generation of transgenic barley lines expressing *R. padi* effectors

Each of the effectors was cloned into the destination vector pBRACT214m containing the ubiquitin promoter from maize for constitutive expression in all plant organs, and a hygromycin marker gene for selection of transgenic lines. Constructs were transformed into the *Agrobacterium* AGL1 strain, supplied with pSOUP, and delivered to the Functional Genomics Facility (FUNGEN) at the James Hutton Institute for *Agrobacterium*-mediated barley embryo transformation of the cultivar Golden Promise. After approximately four months, we obtained different barley lines regenerated from independent calli. The T0 generation was tested for the expression of effector genes by PCR on cDNA from the regenerated plants. RNA was extracted from T0 independent lines using the RNeasy Plant Mini Kit (Qiagen). RNA plant samples were DNAse treated with Ambion® TURBO DNA-free™. SuperScript® III Reverse Transcriptase (Invitrogen) and random primers were used to prepare cDNA. The majority of these plants were positive in PCR tests using effector gene-specific primers (Supplementary Table S1). T1 seeds were germinated on selective media (AgarGel^TM^-containing Hygromycin (100 µg/ml) to select for transformants. Lines showing a 1:3 segregation, representing a single insertion (75 % survival rate on selective media), were selected for further analyses. Universal Probe Library (UPL-Roche Diagnostics ©) was used to quantify effector gene expression in T1 barley transgenic lines ectopically expressing *R. padi* effectors. Barley cv Golden Promise, the background genotype of the transgenic lines, was used as control. RNA from six different barley lines per construct was reverse transcribed into cDNA. Probes and primers (Supplementary Table S1) designed with the UPL System Assay Design (Roche) were tested for at least 95-105% efficiency. Internal controls were Actin-2 (MLOC_78511.2) and Pentatricopeptide (AK373147/MLOC_80089.1) as described previously (Escudero-Martinez *et al*., 2017). Three technical replicates were included for each sample. Relative expression was calculated with the method ΔCt (Delta Cycle threshold) with primer efficiency consideration. One of each of the transgenic effector lines was used as a reference line to calculate the fold-change in additional lines.

Lines positive for effector expression were then bulked into T2 and screened for homozygosity based on complete resistance to hygromycin. Three independent homozygous lines per effector were used to perform the aphid performance assays with *R. padi* and *M. persicae*.

### *M. persicae* performance assays on *N. benthamiana*

Effectors were transiently expressed using vector pB7WG2 in *N. benthamiana* as explained above. The empty vector pB7WG2 was used as a control. Twelve infiltration sites were used per construct per biological replicate (n=12 per biological replicate). One day after infiltration, the abaxial side of the infiltration sites was exposed to two *M. persicae* adults enclosed in a clip cage. The following day, adult aphids were removed leaving three 1^st^ instar nymphs at the underside of the leaves in a clip cage. Seven days later, *N. benthamiana* plants were replaced by freshly infiltrated plants to ensure continued expression of effectors in the plant tissue. After 14 days, the number of nymphs per adult was counted and data were analysed using One way ANOVA (in R-studio) and the post-hoc Fisher’s protected Least Significant Differences (LSD) (cut-off p ≤0.05). Three biological replicates were performed with each replicate containing 12 infiltration sites per construct.

### Aphid performance assays on barley transgenic lines

Seven-day old transgenic barley plants expressing *R. padi* effectors were infested with two 1st instar age-synchronized nymphs (*M. persicae*) or with two 2-day old age-synchronized nymphs (*R. padi*). Barley cv. Golden Promise wild-type plants were used as the control. We used 6-8 plants per individual transgenic line for each biological replicate per aphid species (n=6-8), and four biological replicates were performed. The number of nymphs per adult was monitored at 11 days after infestation for *R. padi*, and after 14 days for *M. persicae*. The resulting data was analysed One-way ANOVA (in R-studio) with post-hoc Fisher’s protected Least Significant Differences (LSD).

### Histochemical GUS staining

To assess GUS expression driven by the maize ubiquitin promoter in transgenic barley transformed with the pBRACT214m-GUS construct we collected different tissues and stained these with 1mg/ml of X-gluc (5-bromo-4-chloro-3-indolyl-B-D-glucuronic acid, Thermo Scientific, USA) in X-gluc buffer (100mM sodium phosphate buffer pH 7.0, 0.1% Triton X-100, 2mM potassium ferricyanide and 2mM potassium ferrocyanide). Tissues were vacuum-infiltrated and incubated in darkness at 37°C overnight. The next day, chlorophyll was removed from the tissues with 1:3 acetic acid/ethanol. Pictures were taken under the dissecting microscope with a Zeiss camera.

### Quantitative RT-PCR to assess defence gene expression in Rp1 transgenic lines

Gene expression of different defence/hormone signalling pathways genes were analysed by qRT-PCR, The transgenic barley Rp1 lines along with the control plants (cv. Golden Promise) were pre-germinated in Petri dishes covered with wet filter paper for three days in the dark at room temperature. Germinated seeds placed on soil and grown under controlled conditions (8h light, 22°C, 70% humidity and 125 µmol photons/m2.s). For basal gene expression the first leaf of the plants (n=6 per genotype) were collected and flash frozen in liquid nitrogen. In addition, barley plants (n=6 plants per transgenic line or wild type control) were exposed to either empty clip cages or clip cages containing 30 mixed-age *R. padi* aphids. Leaf tissues enclosed within the clip cages were collected after 24 h and 72 h. The experiment was performed in three biological replicates (n=6 plants per transgenic or wildtype line per biological replicate) and samples were harvested at the same time of the day: barley plants were treated and collected at 12 am for the 24 h timepoint and at 3 pm for 72 h timepoint, avoiding any effects of the plant circadian cycle.

The local database *Morex genes* was used for retrieving the barely sequences and the *Roche UPL assay design centre* for primer design (Supplementary Table S1). The primers were tested for efficiency (85-115%) and relative gene expression was calculated with the method ΔΔCt (Delta-delta Cycle threshold). Three technical replicates were included for each sample. Cycle threshold values were normalized with two reference genes, pentatricopeptide (AK373147/MLOC_80089) and ubiquitin (AK248472). Expression of these two reference genes was unaffected in our previous microarray experiment (Escudero-Martinez *et al*., 2017). The Wilcoxon Rank Sum Test (cut-off p ≤0.05) was used to assess differences in expression between plant genotypes and treatments.

## Results

### Effector sequence divergence between the aphid species *R. padi* and *M. persicae*

We predicted putative orthologs for 3 previously described *M. persicae* effectors, MpC002, Mp1 and Mp58 from *R. padi* using reciprocal best blast hit analyses on available aphid transcriptome datasets and aphid genome assemblies (threshold of 70 % identity and 50 % query coverage) (Thorpe *et al*., 2016; Thorpe *et al*., 2018). To confirm the sequences of putative orthologous effector pairs we cloned and sequenced their coding sequences.

Amino acid and nucleotide sequence alignments show varying degrees of sequence divergence across the selected effector pairs (Fig. 1, Supplementary Fig. S1). RpC002 is smaller than MpC002, with 193 amino acids compared to 265, respectively, and these effectors share 52.86% sequence identity. The difference in sequence length is partly due to a lack of the NDNQGEE repeat in the N-terminal region of RpC002 (Fig. 1A, Supplementary Fig. S1). Variation in the number of NDNQGEE repeats in MpC002 was previously also detected within *M. persicae* (Thorpe *et al*., 2016), and in this study we characterized the MpC002 version containing 5 repeats. MpC002 also has an extended C-terminal domain compared to RpC002 (Fig. 1A). Rp1, which is similar to *M. persicae* Mp1, is composed of 140 amino acids compared to 139 for Mp1, and these effectors share a percentage sequence identity of 56.12% (Fig. 1B, Supplementary Fig. S1). Lastly, Mp58 and Rp58 contain 152 and 155 amino acids, respectively, share 64.94% sequence identity, and are most divergent in the C-terminal region of the protein (Fig. 1C, Supplementary Fig. S1).

**Fig. 1.**
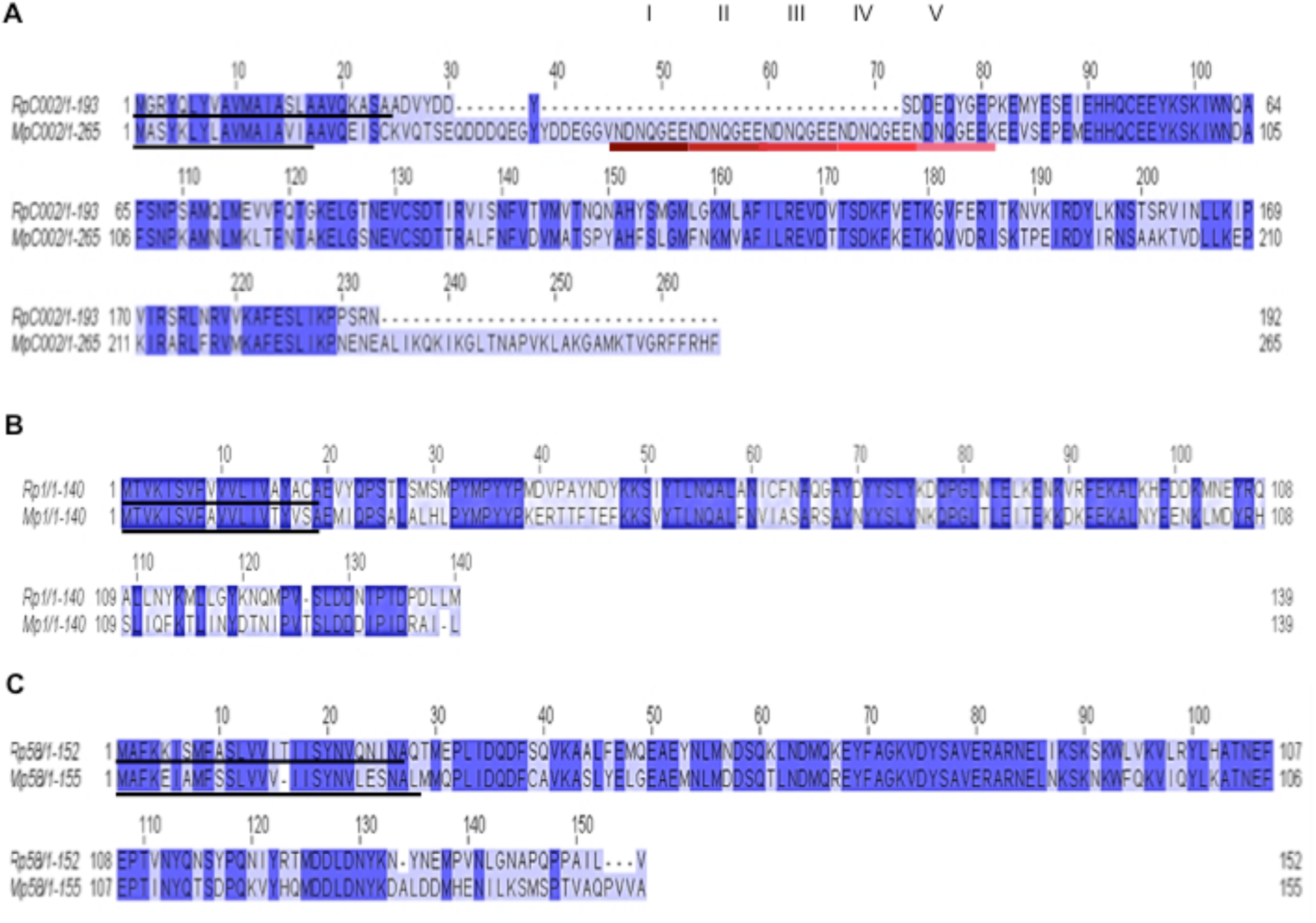
Pair-wise amino acid sequence alignments of three selected effectors from *Rhopalosiphum padi* and *Myzus persicae*. Alignments were generated using Jalview 2.10.4. The level of sequence conservation is indicated by dark (high identity) to light purple colour (low identity). Predicted signal peptide (Signal P4.1) sequences are underlined in black. **A**) RpC002/MpC002 alignment. The 5x repeat motif (NDNQGEE) in MpC002 is underlined with different shades of red to pink. **B**) Rp1/Mp1 alignment. **C**) Rp58/Mp58 alignment.

### Effector gene expression is consistent across different feeding/plant environments, but the range of expression varies between aphid species

We were interested in assessing how gene expression of the three effector pairs was affected in *R. padi* and *M. persicae* upon exposure to different feeding/plant environments. We made use of previously generated aphid RNAseq datasets (Thorpe *et al*., 2018) to investigate gene expression of our effectors of interest by plotting their gene counts across different treatments (exposure to diet, host and poor/nonhost plants) and timepoints (3h and 24h exposure). All 6 aphid effectors were expressed with only limited variation in expression across the different aphid treatments and timepoints (Fig. 2). Whilst the three selected effectors from *M. persicae* displayed more similar gene expression levels compared to one another, ranging from 280 counts for *MpC002* to 904 counts for *Mp1* (Fig. 2B and C), the three effectors from *R. padi* showed a wider range of expression. For instance, gene counts varied from 271 for *RpC002* to 2112 for *Rp1* over the various treatments and timepoints (Fig. 2A).

**Fig. 2.**
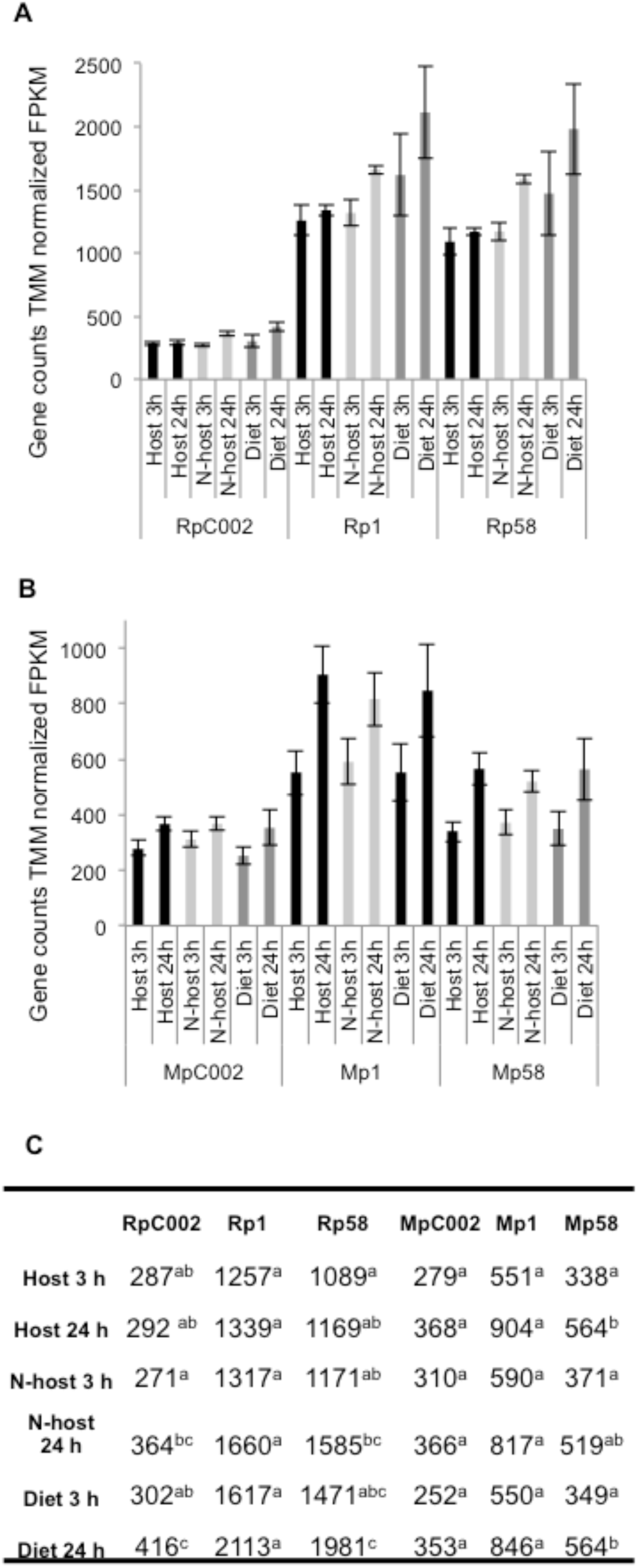
Effector gene expression in *Rhopalosiphum padi* and Myzus persicae upon exposure to different feeding environments. **A**) Expression of *R. padi* effectors *RpC002, Rp1* and *Rp58* upon aphid exposure to barley (host), Arabidopsis (non-host) or artificial diet for 3h or 24h. The expression of transcripts was normalized by the TMM-FPKM method (fragments per kilo-base of exon per million reads mapped). Bars indicate standard error. **B**) Expression of *M. persicae* effectors *MpC002*, *Mp1* and *Mp58* upon aphid exposure to Arabidopsis (host), barley (poor-host) or artificial diet. The expression of transcripts was normalized by the TMM-FPKM method (fragments per kilo-base of exon per million reads mapped). Bars indicate standard error. **C**) Table displaying expression values for each effector. Letters indicate significant differences as determined by one-way ANOVA and post-hoc protected Least Significant Differences (p<0.05 *; p<0.01 ***).

### *R. padi* effectors show similar subcellular localisation as their putative *M. persicae* orthologs *in N. benthamiana*

The subcellular localization of effectors can provide important information on the cellular compartment that is targeted by these proteins. We used confocal microscopy of GFP-tagged *R. padi* effectors alongside their *M. persicae* putative orthologs to compare subcellular localisation *in planta*. The GFP-effector fusion proteins (N-terminal GFP tag) were transiently expressed in leaves of *N. benthamiana*, which is a host for *M. persicae*, but a non-host for *R. padi*. Western blotting showed that all GFP-fusion proteins were expressed, but that two of the *R. padi* effectors, RpC002 and Rp58, showed lower protein levels than their putative *M. persicae* orthologs (Supplementary Fig. S2), with RpC002 only detected once in three biological replicates (Supplementary Fig. S2). In contrast, Rp1 from *R. padi* was detected more strongly than its putative *M. persicae* ortholog Mp1 (Supplementary Fig. S2). We detected GFP signal corresponding to the RpC002- and MpC002-fusion proteins by confocal microscopy at the plasma membrane, and in some cases, a weak signal was present in the nucleus or the cytoplasm (Fig. 3A and Supplementary Fig. S3). It should be noted that expression of RpC002 was very low, especially compared to MpC002 (Supplementary Fig. S2), and only a few transformed cells were visible. We validated the plasma membrane localization of RpC002/MpC002 effectors upon co-expression with a plasma membrane marker (Nelson *et al*., 2007) (Fig. 3A). Both the Rp1 and Rp58 were detected in the cytoplasm and nucleus, similar to their putative *M. persicae* orthologs and the free GFP control (Fig. 3B and Fig. 3C). Similarly, we tried to express tagged effectors in barley epidermal cells using particle bombardment, but due to low signal we were unable to reliably localize effectors in this system. Overall, the three selected *R. padi* effector showed similar subcellular localization patterns in *N. benthamiana* as their putative *M. persicae* orthologs.

**Fig. 3.**
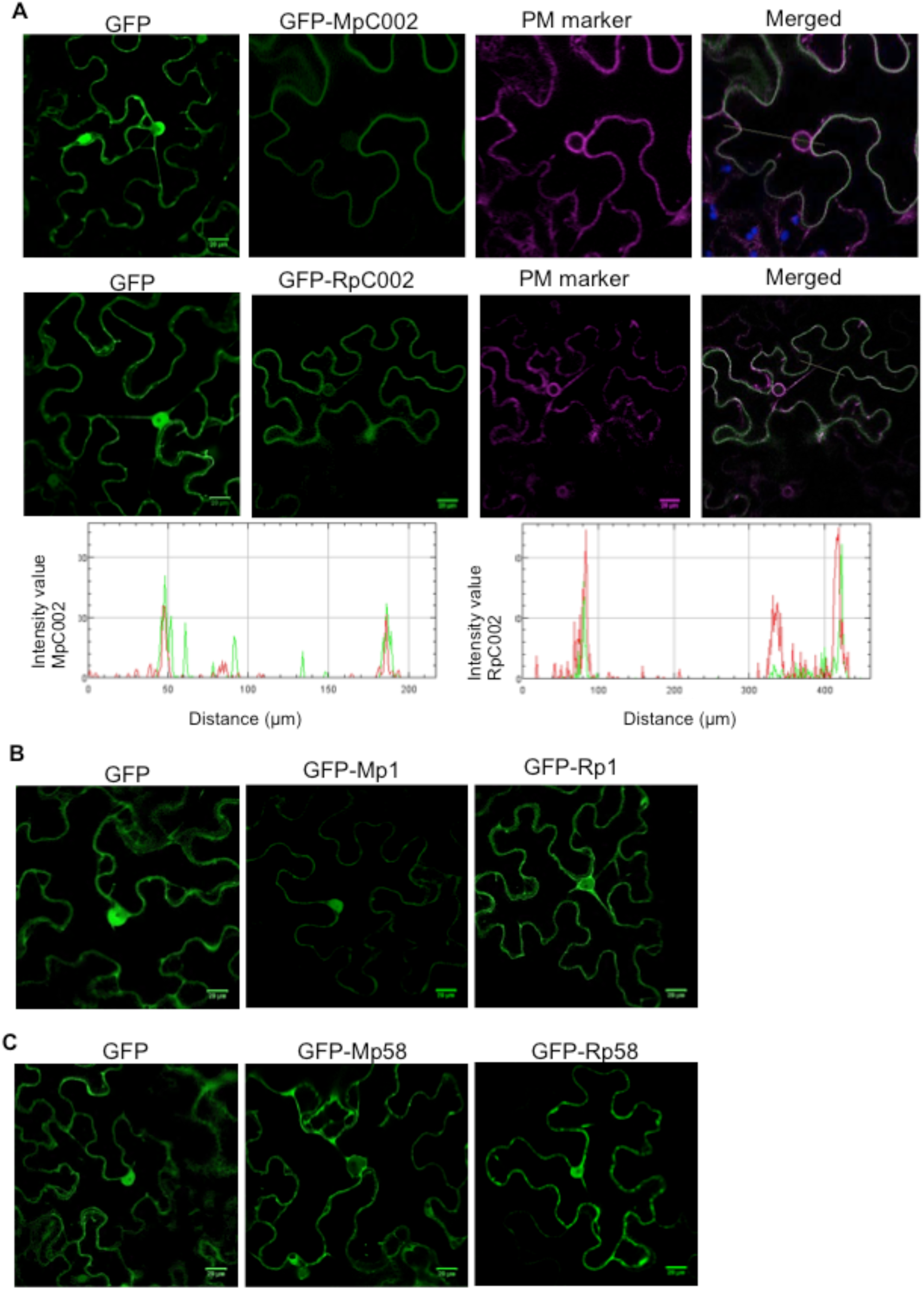
Localization of aphid effectors in Nicotiana benthamiana. **A**) Confocal microscopy images of free GFP (empty vector, pB7WG2F), and effectors GFP-MpC002 and GFP-RpC002 (middle section) transiently overexpressed in *N. benthamiana*. Both effectors were co-expressed with a plasma membrane marker (PM marker; Nelson et al., 2007). Merged pictures represent the overlay image of the GFP and RFP channels. Graphs represent overlay between the fluorescence intensity of the GFP (green series) and RFP (red series) channels of a particular selected region of interest (ROI) shown as a line in the merged picture. The images were taken 3 days after agroinfiltration. The co-localization was analysed by Fiji software and the plugin RGB profiler. **B**) Confocal microscopy images of free GFP alone (pB7WG2F), and effectors GFP-Mp1 / GFP-Rp1. **C**) Confocal microscopy images of free GFP alone (pB7WG2F), and effector GFP-Mp58 / Rp58.

### Expression of Rp58 in *Nicotiana benthamiana* reduces host susceptibility to *M. persiae*

To assess whether the three selected *R. padi* effectors can impact host susceptibility to *M. persicae* when expressed in a *R. padi* nonhost plant species, we performed aphid performance assays on *N. benthamiana* leaves transiently expressing the different effectors under the control of a 35S promoter. In line with previous reports (Bos *et al*., 2010) (Pitino and Hogenhout, 2013), we found that ectopic expression of MpC002 significantly increased the number of *M. persicae* nymphs produced per adult by 27% (One-way ANOVA post-hoc Fisher’s protected Least Significant Differences (LSD); p>0,05) (Fig. 4A). In contrast, RpC002 did not alter *N. benthamiana* host susceptibility to *M. persicae*. Western blot analyses of GFP-MpC002 and GFP-RpC002 showed that RpC002 protein is detected at a much lower level than MpC002 (Supplementary Fig. S2), and therefore it is possible that also the untagged MpC002 and RpC002 proteins expressed in the aphid performance assays have different levels of abundance which affects the phenotypic observations. No significant differences in host susceptibility were noted upon expression of Mp1 from *M. persicae* and Rp1 from *R. padi* (Fig. 4B), in line also with a previous report that transient expression of Mp1 under the 35S promoter in *N. benthamiana* does not affect susceptibility (Bos *et al*., 2010). The expression of Mp58, but not Rp58, resulted in significantly lower *M. persicae* nymph production compared to the vector control, with 55% and 27% less nymphs being produced per adult, respectively (One way ANOVA post-hoc Fisher’s protected Least Significant Differences (LSD); p>0,05) (Fig. 4C).

**Fig. 4.**
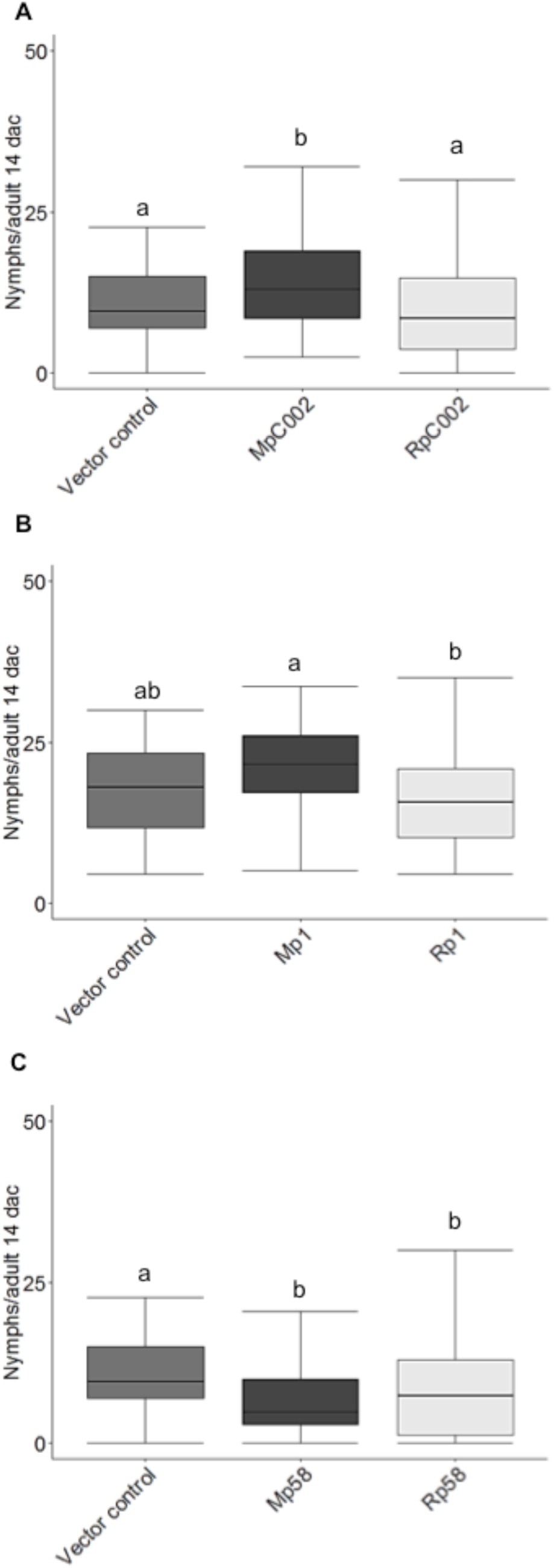
*Myzus persicae* performance on *Nicotiana benthamiana* plants expressing aphid effectors. Leaves of *N. benthamiana* were agroinfiltrated with different effector constructs (35S-promoter) and infiltration sites were challenged with 3 *M. persicae* nymphs, which were allowed to develop and reproduce. Nymph production per aphid was monitored over a 14-day period, with the aphids being moved to freshly infiltrated leaves every 7 days. Empty vector was used as a control. **A**) Number of nymphs produced per adult on *N. benthamiana* leaves expressing the vector control, MpC002 or RpC002. **B**) Number of nymphs produced per adult on *N. benthamiana* leaves expressing the vector control, Mp1 or Rp1. **C**) Number of nymphs produced per adult on *N. benthamiana* leaves expressing the vector control, Mp58 or Rp58. Box plots show the average number of nymphs per adult 14 days after challenge (dac) from three independent biological replicates (number of plants per effector or control used on each replicate = 12). Different letters indicates significant differences at p>0.05. Statistical analyses were performed using one-way ANOVA post-hoc Fisher’s protected Least Significant Differences (p>0.05).

### Expression of RpC002 and Rp1 in transgenic barley enhances susceptibility to *R. padi*

Aphid effector characterization studies to date have focused on dicot plant species. With *R. padi* being a major pest of cereals, we aimed to extend aphid effector characterization studies to the monocot crop barley to explore the contribution of *R. padi* effectors to host susceptibility. We generated barley transgenic lines in the cultivar Golden Promise to ectopically express the three *R. padi* effectors Rp1, RpC002 and Rp58 using a modified version of the pBRACT214 vector (Colas *et al*., 2019), containing the ubiquitin promoter from maize to allow constitutive expression in all plant organs (http://www.bract.org/constructs.htm#barley). To determine where candidate genes of interest are potentially expressed when transformed into barley using this pBRACT214m construct, we performed GUS-staining on different plant tissues, such as leaves, stems, spikes and roots of a barley transgenic line generated by transformation with pBRACT214m:GUS (Supplementary Fig. S4). We observed GUS expression in all tissues, with particularly strong expression in leaf vascular tissues and roots (Supplementary Fig. S4). After barley transformation, we obtained 13 independent lines for the RpC002 effector, 4 lines for the Rp1 effector, and 16 lines for Rp58. In the first generation, lines with a single effector insertion were selected based on around 75% survival on Hygromycin (hygromycin phosphotransferase is the pBRACT selection marker), yielding 8 independent transgenic lines for RpC002, 3 lines for Rp1, and 7 lines for Rp58. The presence of effector coding sequences (lacking the signal peptide encoding sequence) was confirmed in the T0 generation by semi quantitative RT-PCR (data not shown) and was verified in the T1 generation by qRT-PCR (Supplementary Fig. S5). We did not observe any visual differences in plant growth and development for any of the transgenic lines selected (Supplementary Fig. S5). Three homozygous T3 lines per effector construct were selected for aphid performance assays with *R. padi* and *M. persicae* to assess how barley host and poor-host interactions with aphids were affected. Each plant was infested with two nymphs and reproduction was assessed after 11 days for *R. padi* and after 14 days for *M. persicae*.

For *M. persicae* we did not find consistent significant differences in aphid performance on the barley transgenic lines expressing the *R. padi* effectors compared to the wild-type control (Fig. 5). For one of the Rp1 lines, Rp1_2A, however, we noted increased nymph production (Fig. 5B). Line Rp1_2A was also the highest Rp1 expressing line we identified (Supplementary Fig. S5), and additional lines with similar expression levels would need to be identified to rule out the possibility that the insertion site in this line causes the enhanced susceptibility phenotype.

**Fig. 5.**
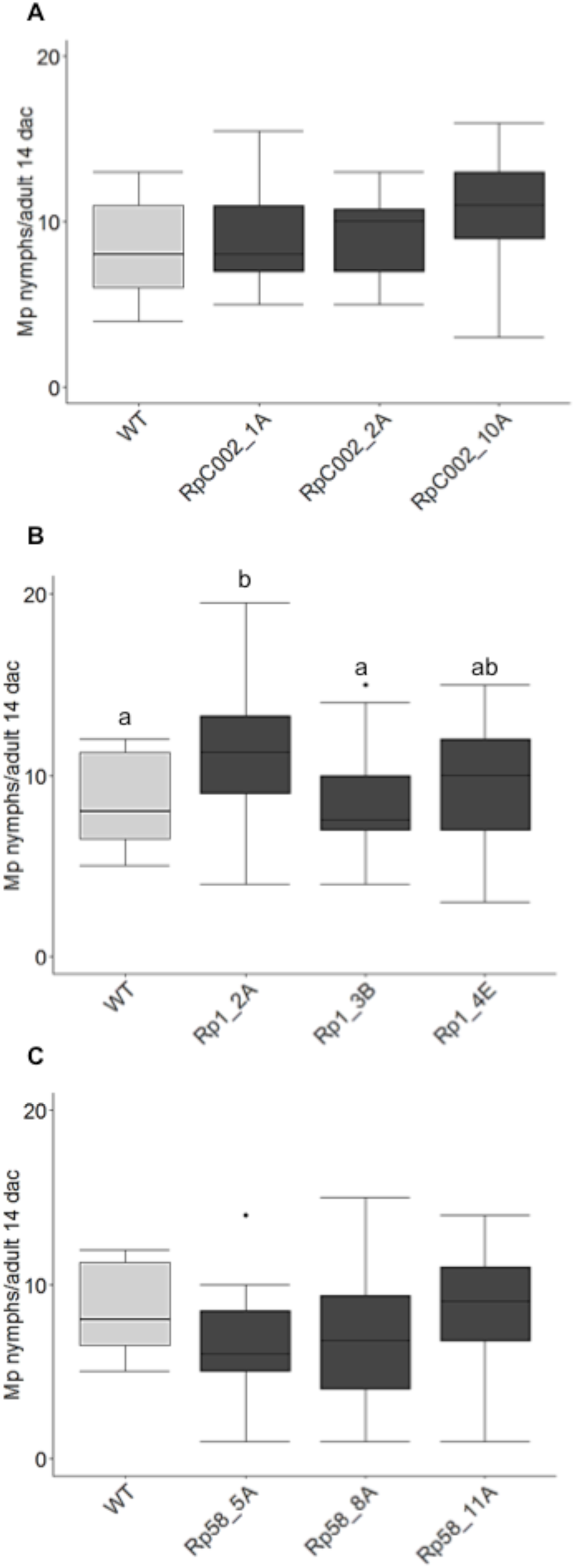
Myzus persicae performance on barley plants ectopically expressing different *Rhopalosiphum padi* effectors. Transgenic barley lines were challenged with aphids alongside wild-type plants cv Golden Promise (WT). Nymph production was monitored for 14 days. **A**) Nymph production per adult on transgenic barley lines expressing effector RpC002. Three independent transgenic lines were assessed: RpC002_1A, RpC002_2A and RpC002_10A. **B**) Nymph production per adult on transgenic barley lines expressing effector Rp1. Three independent transgenic lines were assessed: Rp1_2A, Rp1_3B and Rp1_4E. **C**) Nymph production per adult on transgenic barley lines expressing effector Rp58. Three independent transgenic lines were assessed: Rp58_5A, Rp58_8A and Rp58_11A. Box plots show the average number of nymphs per adult 14 days after challenge (dac) from at least three independent biological replicates (number of plants per effector or control used on each replicate = 6-8). Different letters indicate significant differences as determined with one-way ANOVA post-hoc Fisher’s protected least significant difference test (p>0.05).

Ectopic expression of both RpC002 and Rp1 in transgenic barley lines enhanced susceptibility to *R. padi* (Fig. 6A and B). Specifically, two out of three independent RpC002 barley lines, RpC002_1A and RpC002_2A, showed 16% and 12% increased nymph production compared to the wild-type control, respectively (One way ANOVA post-hoc Fisher’s protected Least Significant Differences (LSD); p>0,05) (Fig. 6A). The transgenic line with the strongest susceptibility phenotype (RpC002_1A) also showed the highest *RpC002* expression level (Fig. 6A; Supplementary Fig. S5). In addition, all three independent Rp1 barley lines showed enhanced susceptibility to *R. padi* with an increased nymph production of 11-22% across lines compared to the wild-type control (One way ANOVA post-hoc Fisher’s protected Least Significant Differences (LSD); p>0,05) (Fig. 6B). Also, we observed a correlation between the level of effector gene expression and impact on host susceptibility to aphids, with the lines showing the most pronounced susceptibility phenotype towards *R. padi* (Rp1_2A and Rp1_3B) also showing the higher *Rp1* transcript levels (Fig. 6B; Supplementary Fig. S5).

**Fig. 6.**
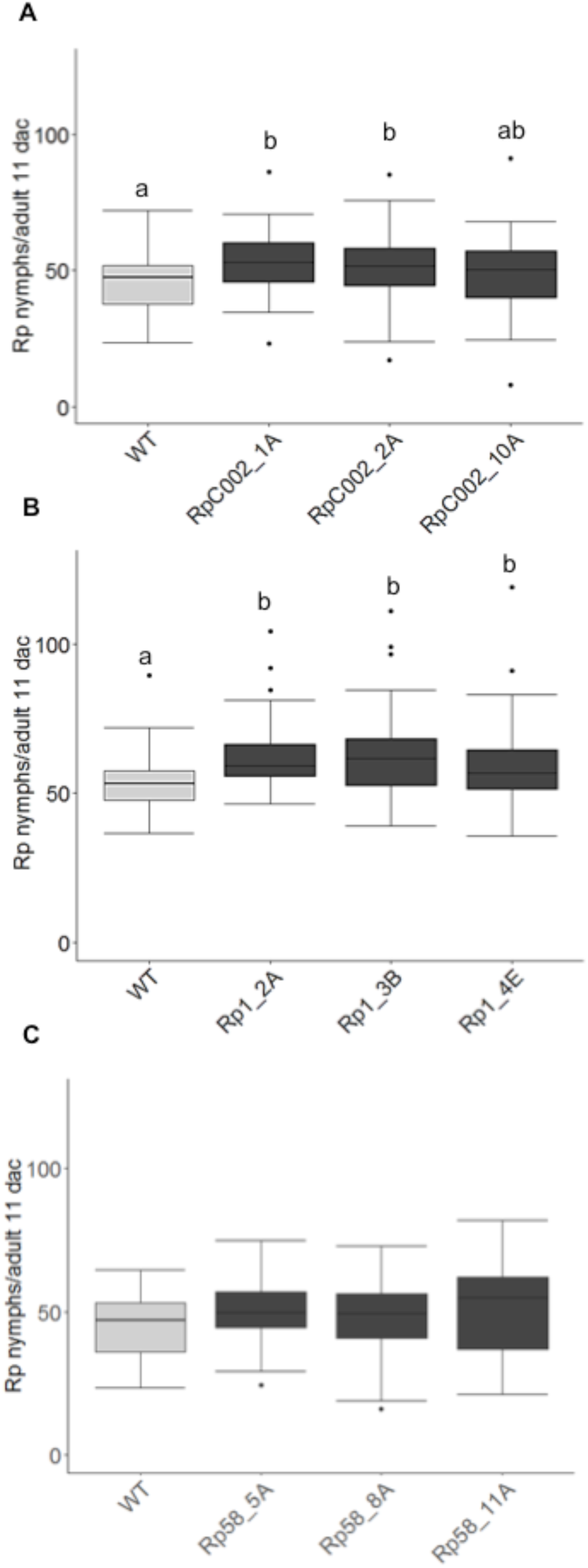
*Rhopalosiphum padi* performance on barley plants ectopically expressing different *R. padi* effectors. Transgenic barley lines were challenged with aphids alongside wild-type plants cv Golden Promise (WT). Nymph production was monitored for 11 days. **A**) Nymph production per adult on transgenic barley lines expressing effector RpC002. Three independent transgenic lines were assessed: RpC002_1A, RpC002_2A and RpC002_10A. **B**) Nymph production per adult on transgenic barley lines expressing effector Rp1. Three independent transgenic lines were assessed: Rp1_2A, Rp1_3B and Rp1_4E. **C**) Nymph production per adult on transgenic barley lines expressing effector Rp58. Three independent transgenic lines were assessed: Rp58_5A, Rp58_8A and Rp58_11A. Box plots show the average number of nymphs per adult 11 days after challenge (dac) from at least three independent biological replicates (number of plants per effector or control used on each replicate = 5-10). Different letters indicate significant differences as determined with one-way ANOVA post-hoc Fisher’s protected least significant difference test (p>0.05).

No differences were observed for the Rp58 transgenic lines, which showed susceptibility levels similar to the wild-type control.

### Rp1 suppresses defence signalling in transgenic barley lines

To gain further insight into how Rp1 may enhance barley host susceptibility to aphids, we investigated the basal and induced expression levels of defence-related genes in the three independent transgenic lines we generated (lines 2A, 3B and 4E), compared to the wild-type cultivar Golden Promise. Based on our previous work (Escudero-Martinez *et al*., 2017) we selected barley genes strongly induced upon *R. padi* infestation: *beta-thionin* (AK252675), *SAG12-like* (MLOC_74627.1), a *jasmonate ZIM domain gene 3* (*JAZ3*, MLOC_9995), *lipoxygenase 2* (*LOX2*, MLOC_AK357253) and the *jasmonate-induced gene* (*JI*, MLOC_15761). We further expand our selected genes set based on markers of different hormone signalling pathways, with focus on the jasmonate pathway, which is strongly activated upon aphid infestation (Escudero-Martinez *et al*., 2017): *lipoxygenase 5* (*LOX5*, MLOC_71948), the *WRKY transcription factor 50* (*WRKY50*, MLOC_66204), the *allene cyclase oxydase* (*AOC*, MLOC_68361) and *jasmonate-induced gene 2* (*JI2*, MLOC_56924) markers for jasmonate; but also the salicylic acid marker *non-expressor of pathogenesis-related 1-like* (*NPR1*, AM050559.1), *the ethylene-response factor 1* (*EFR1*, MLOC_38561) and *abscisic acid-inducible late embryogenesis abundant 1* (*A1*, MLOC_72442). We analysed basal gene expression levels as well as expression levels upon 24h and 72h exposure to clip cages with or without aphids. It should be noted that the use of clip-cages, even when empty, triggers changes in gene expression due to mechanical damage, and that all selected genes were induced by aphid challenge (Supplementary Fig. S6 and S7).

First, we compared basal gene expression levels across plant lines not infested with aphids and without being exposed to a clip-cage (Fig. 7A). We found that expression of a gene encoding a SAG-12 like cysteine protease (MLOC_74627.1) was most strongly reduced at basal levels in Rp1 lines compared to the wild-type control, but differences were only significant for lines Rp1-2A and Rp1-4E, possibly due to sample variation for line Rp1-3B (Fig. 7A). *SAG12-like* expression was also reduced in the transgenic lines upon exposure to either empty clip cages (Supplementary Fig. S6A and B) or clip cages containing aphids (Fig. 7B and C), compared to wild-type plants, but not consistently to statistically significant levels. The basal level of *EFR1* expression was slightly but significantly higher in Rp1 transgenic lines compared to the wild-type control (Fig. 7A).

**Fig. 7.**
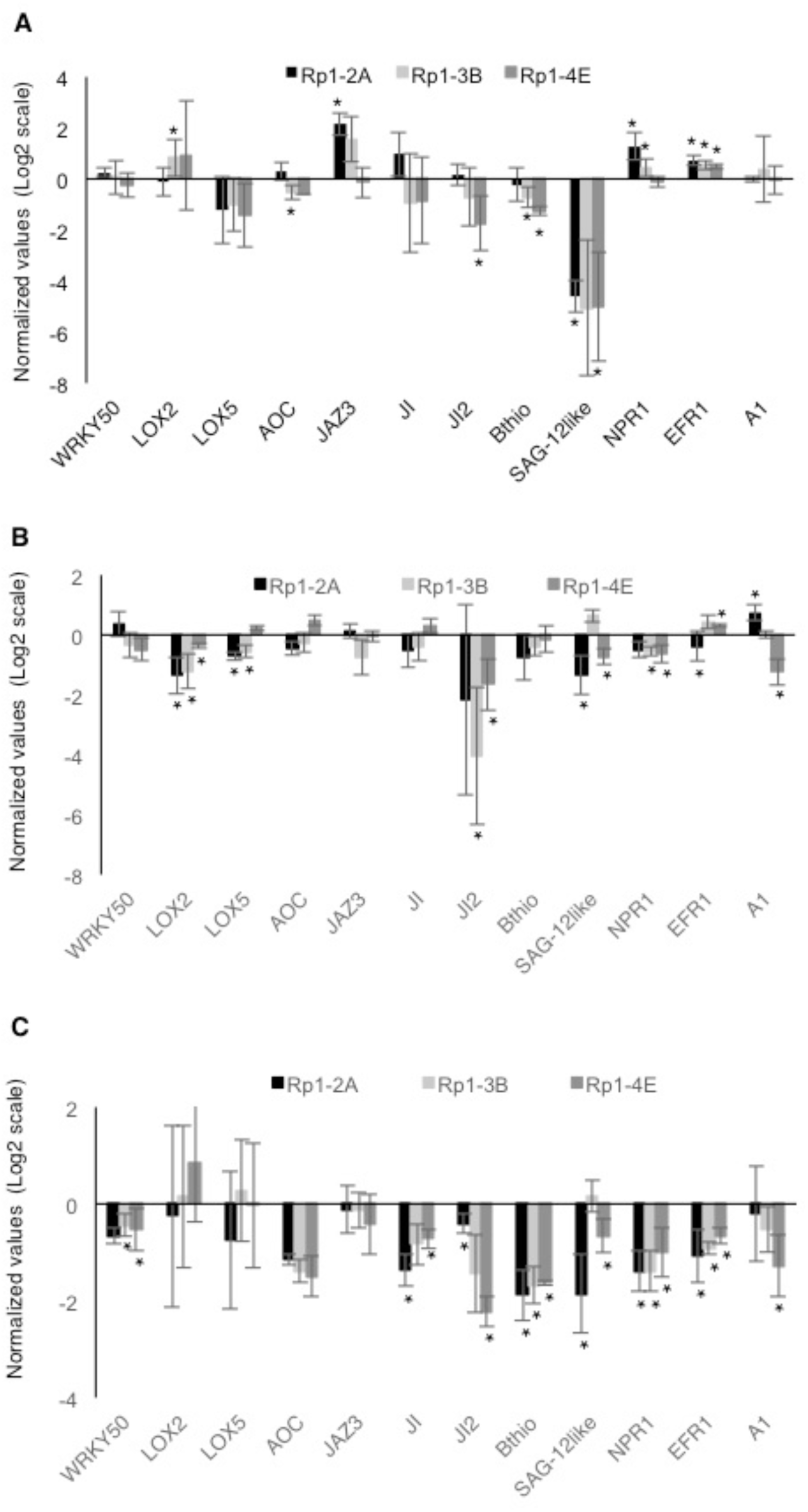
Basal and aphid-induced defence gene expression in barley Rp1 lines. Relative gene expression of defence-related/hormone-signalling genes was measured by qRT-PCR in control barley plants (cv. Golden Promise) and three independent barley lines expressing the R. padi effector Rp1. **A**) Log-fold changes of barley basal gene expression (no aphids, no clip cage) in three transgenic Rp1 barley lines relative to control lines (WT=0). **B**) Log-fold changes of barley gene expression upon 24h exposure to clip cages with R. padi aphids in three transgenic Rp1 barley lines relative to control plants (WT=0). **C**) Log-fold changes of barley gene expression upon 72h exposure to clip cages with R. padi aphids in three transgenic Rp1 barley lines relative to control lines (WT=0). All gene expression analyses were based on three independent biological replicates, and graphs represent mean expression. Genes are represented in the graphs are: WRKY transcription factor 50 (WRKY50, MLOC_66204), lipoxygenase 2 (LOX2, MLOC_AK357253), lipoxygenase 5 (LOX5, MLOC_71948), allene cyclase oxydase (AOC, MLOC_68361), jasmonate ZIM domain gene 3 (JAZ3, MLOC_9995), jasmonate-induced gene (JI, MLOC_15761), jasmonate-induced gene 2 (JI2, MLOC_56924), beta-thionin (AK252675), SAG12-like (MLOC_74627.1), non-expressor of pathogenesis-related 1-like (NPR1, AM050559.1), the ethylene-response factor 1 (EFR1, MLOC_38561) and abscisic acid-inducible late embryogenesis abundant 1 (A1, MLOC_72442). Black bars represent gene expression levels in Rp1-2A lines, light grey bars represent gene expression levels in Rp1-3B lines and grey bars gene expression levels in Rp1-4E lines. Asterisks indicate significant differences between control plants (WT) and Rp1 transgenic lines (Wilcoxon Rank Sum Test, p ≤0.05).

In addition, four genes (*WRKY50*, *AOC*, *beta-thionin*, *NPR1*) were significantly less expressed across all transgenic lines compared to the wild-type control when leaves were exposed for 24h to empty clip-cages (Supplementary Fig. S6A and B). *LOX2*, *JI*, and *JI2* showed a trend towards reduced expression in transgenic lines, but differences were not consistently significant across all lines (Supplementary Fig. S6A and B). In response to clip-cages with aphids for 24h, only *LOX2* showed a significant reduction in expression in all transgenic lines, whereas *JI2* reduced expression was noticeable not consistently significant (Fig. 7B). For the 72h timepoint, 3 marker genes (*beta-thionin*, *NPR1*, *EFR1*) showed a significant reduction in expression in all lines compared to wild-type plants when exposed to clip-cages containing aphids, and similar trends were observed for *WRKY50*, *JI*, *JI2* and *SAG12-like* (Fig. 7C). Overall, we observed a reduction of several marker genes of defence/hormone signalling pathways relevant to plant-aphid interactions in the Rp1 transgenic barley lines, which may translate into their enhanced susceptibility to aphids.

## Discussion

Aphids are damaging pests on cereals, including barley. Aphid effector characterization efforts to date have focused on dicot plant species including Arabidopsis, tomato, and *Nicotiana benthamiana* (Mutti *et al*., 2008) (Bos *et al*., 2010) (Pitino *et al*., 2011) (Atamian *et al*., 2013) (Rodriguez *et al*., 2014) (Elzinga *et al*., 2014; Pitino and Hogenhout, 2013; Rodriguez *et al*., 2017a) (Chaudhary *et al*., 2019), and have not yet been described for monocot crops. It is crucial to understand the mechanisms employed by aphids and other insects to infest cereals, as well as to gain insight into how aphid effector function may have diverged across different plant-aphid species interactions. Although challenging, functional characterization of aphid effectors not only in dicot (model) plants, but also in monocot crops, promises to reveal novel insight into effector function and evolution.

Effector diversity across different aphid species might reflect the adaptation to different host plants (Schulze-Lefert and Panstruga, 2011). Amino acid alignments of the putative orthologous effectors we selected showed different levels of sequence divergence which might reflect the different lifestyles of the two aphid species *R. padi* (cereal specialist) and *persicae* (broad host range pest). In general, the signal peptide sequences of these effectors tend to be more conserved than their C-terminal regions, indicating divergence mainly occurred within the functional effector domains. The NDNQGEE repeat motif, which is absent in RpC002, was previously shown to be linked to virulence in *M. persicae*, since MpC002 transgenic Arabidopsis lines, but not lines expressing a deletion mutant missing the repeat motifs, showed enhanced susceptibility to aphids (Pitino and Hogenhout, 2013). We noticed that the RpC002 protein, which lacks the NDNQGEE repeats, is less expressed/stable in *N. benthamiana*, which could explain the limited impact on plant susceptibility in this plant species. Noteworthy, within *M. persicae*, different MpC002 variants have been reported with different numbers of the NDNQGEE repeat (Thorpe *et al*., 2016). The biological significance of this repeat variation remains to be elucidated.

All selected aphid effectors were expressed regardless whether aphids were exposed to a host, non-host plant or artificial diet. It is possible that, unlike the case for plant pathogens where effector gene expression varies across different infections stages (Cotton *et al*., 2014; Hacquard *et al*., 2013; Jupe *et al*., 2013; O’Connell *et al*., 2012), aphid effectors are constitutively expressed to ensure aphids are generally ready to infest a plant. This hypothesis is in line with other reports where no significant overall effector gene expression variation was reported when aphids were adapted to different plant environments (Lu *et al*., 2016; Mathers *et al*., 2017; Thorpe *et al*., 2018). The Rp1/Mp1 and Rp58/Mp58 pair was more similarly expressed in the two aphid species relative to the RpC002/MpC002 effector pair. Interestingly, the Mp1- and Mp58-like effectors are co-located in a non-syntenic region across the genomes of 5 different aphid species, and their expression is tightly co-regulated with a large set of aphid genes, including many (predicted) effectors such as MpC002 (Thorpe *et al*., 2018). Whether and how these effectors work together to enable aphid infestation remains to be explored.

Both RpC002 from *R. padi* and MpC002 from *M. persicae* showed similar localisation to the plasma membrane in *N. benthamiana* indicating this could be their sites of activity. Both the nucleus and plant plasma membrane play key roles in activating plant defences against plant pathogenic microbes (reviewed by (Boutrot and Zipfel, 2017; Motion *et al*., 2015)). The plasma membrane is the site of many immune receptors, such as receptor-like kinases, required for pathogen recognition and initiation of an immune response (Boutrot and Zipfel, 2017). The plasma membrane localization of these aphid effectors might reflect a role in interfering with immune receptors or any other cell membrane associated defences. It should be noted that effector localization using highly expressed effectors (35S-based) may be affected by (endogenous) expression levels of their host targets. For example, in the case where the host target is in low abundance in the epidermal cells analysed by confocal microscopy, the highly abundant effector may not visibly localize to the site of the endogenous target. This is the case for Mp1, which only co-localizes to vesicles in the presence of over-expressed VPS52 (interacting host protein), with endogenous levels of VPS52 being low in leaf tissues (Rodriguez *et al*., 2017a).

Despite a similar subcellular localization of MpC002 and RpC002 in *N. benthamiana*, only MpC002 enhanced plant susceptibility in this plant species to *M. persicae*. Species-specific activity within the aphid C002 family was previously reported and linked the presence/absence of the NDNQGEE repeat motif (Pitino and Hogenhout, 2013). In contrast, RpC002 increases barley susceptibility to *R. padi*, indicating the effector is functional when expressed in an appropriate host plant. Whether the NDNQEE motif is associated with reduced protein expression and/or stability in certain plant species remains to be investigated.

Both Rp58 and Mp58 similarly reduced *N. benthamiana* susceptibility to *M. persicae* pointing to a potentially conserved function of these effectors. The reduction in susceptibility mediated by Mp58 is in line with a report by Elzinga et al., (Elzinga *et al*., 2014). Potentially, the artificially high levels of Rp58/Mp58 expression leads to an exaggerated host targeting response and subsequent activation of defences. Alternatively, Rp58/Mp58 was not expressed in tissues where these effectors are usually delivered and active, or these proteins may only function in combination with additional effectors in enhancing plant susceptibility. In contrast to our observations, Atamian et al., (Atamian *et al*., 2013) reported that the putative ortholog of Rp58/Mp58 in *Macrosiphum euphorbiae* (Me10) increased tomato and *N. benthamiana* susceptibility to the potato aphid. Perhaps these effectors function in a different way across plant-aphid interactions.

The lack of an impact of Rp1 and Mp1 on *N. benthamiana* susceptibility to *M. persicae* was not surprising as it was previously shown that Mp1, when expressed under the 35S promoter, does not alter plant susceptibility (Bos *et al*., 2010) (Elzinga *et al*., 2014). However, when expressed under a phloem-specific promoter, Mp1, but not Rp1, enhances *benthamiana* susceptibility to *M. persicae* (Pitino and Hogenhout, 2013) (Rodriguez *et al*., 2017a). Interestingly, Rp1 expression in barley, driven by a ubiquitin-promoter, enhanced barley susceptibility to *R. p*adi but not to the same extent to *M. persicae*, suggesting that not only Mp1, but also Rp1, promotes aphid susceptibility in a specific plant-aphid system. In contrast to the 35S promoter in *N. benthamiana*, the maize ubiquitin promoter in transgenic barley lines is expressed quite strongly in the plant vasculature, which likely is the plant tissue where many aphid effectors function. Barley resistance to *M. persicae* is likely phloem-based (Escudero-Martinez *et al*., 2019) and barley transcriptional responses to this aphid species include a strong activation of a specific set of defence-related genes (Escudero-Martinez *et al*., 2017). It is possible that effectors from the cereal specialist *R. padi* do not affect barley resistance mechanisms against *M. persicae* and as a result susceptibility remains comparable to wild-type plants.

The effect of Rp1 on barley susceptibility to *R. padi* is likely associated with the suppression of several defence genes we observed in transgenic lines expressing this effector. *SAG12-like* encodes a cysteine protease involved in hypersenescence and has been implicated in Arabidopsis PAD4-mediated defence against aphids (Pegadaraju *et al*., 2007). Barley genes encoding beta-thionins contribute to defence against aphids (Escudero-Martinez *et al*., 2017), as well as encoding for components of the JA signaling pathway (reviewed by (Züst and Agrawal, 2016). For example, *LOX2* overexpression in barley increased resistance towards *R. padi* and *M. persicae*, possibly by activating a group of JA-related genes. In line with this, knock-down of *LOX2* in barley resulted in enhanced susceptibility to these same aphid species (Losvik *et al*., 2017). The WRKY50 transcription factor also is implicated in JA signalling, but negatively regulates JA responses while promoting SA-induced expression of PR1 (Gao *et al*., 2011; Hussain *et al*., 2018). The mechanical damage caused by the clip cages may have led to activation of WRKY50, at reduced levels in Rp1 transgenic lines compared to the control. The consistent reduction of both SA and ethylene signalling markers (*NPR1* and *EFR1*) 72h after aphid exposure in the Rp1 transgenic, despite higher basal levels in most of the lines, suggests that defence pathways are suppressed upon expression of the Rp1 effector. Our work represents an important step towards understanding the function of aphid effectors promoting susceptibility in a monocot crop. The future identification of barley host targets of effectors like Rp1, will help us further link the observed suppression of defence gene expression to host susceptibility and reveal the underlying mechanisms.

## Supplementary data

Fig. S1. Pair-wise nucleotide sequence alignments of putative orthologous effectors from *Rhopalosiphum padi* and *Myzus persicae*.

Fig. S2. Western blots showing the expression of GFP and the GFP-effector fusion proteins in *Nicotiana benthamiana*.

Fig. S3. Localization of *Rhopalosiphum padi* effector RpC002 in *Nicotiana benthamiana*.

Fig. S4. Expression of GUS (β-glucuronidase) under control of the maize ubiquitin promoter in different tissue of the barley transgenic line generated using pBRACT214:GUS.

Fig. S5. Effector transcript levels in transgenic barley lines expressing *Rhopalosiphum padi* effectors and plant phenotypes.

Fig. S6. Defence-related gene expression in barley Rp1 lines after exposure to empty clip cages without *Rhopalosiphum padi*.

## Acknowledgements

We thank Abdellah Barakate (JHI, UK) for help and advice on selection of barley transgenic lines and for providing the pBRACT214m construct, Peter Thorpe for help and advice on analysing gene expression of effectors across RNAseq datasets.

This work was supported by the Biotechnology and Biological Sciences Research Council (grant no. BB/J005258/1 to J.I.B.B.), the European Research Council (grant no. APHIDHOST-310190 to J.I.B.B.

## Author contributions

J.I.B.B. conceived and directed the project; C.E-M and J.I.B.B designed the experiments; C.E-M, P.A.R. and P.A.S performed the experiments; C.E-M, P.A.R. and J.I.B.B. analysed the data; J.S. generated the barley transgenic lines; C.E-M and J.I.B.B. wrote the manuscript; all authors approved the final manuscript.

