## Supplementary data for "An aphid effector promotes barley susceptibility through suppression of defence gene expression"

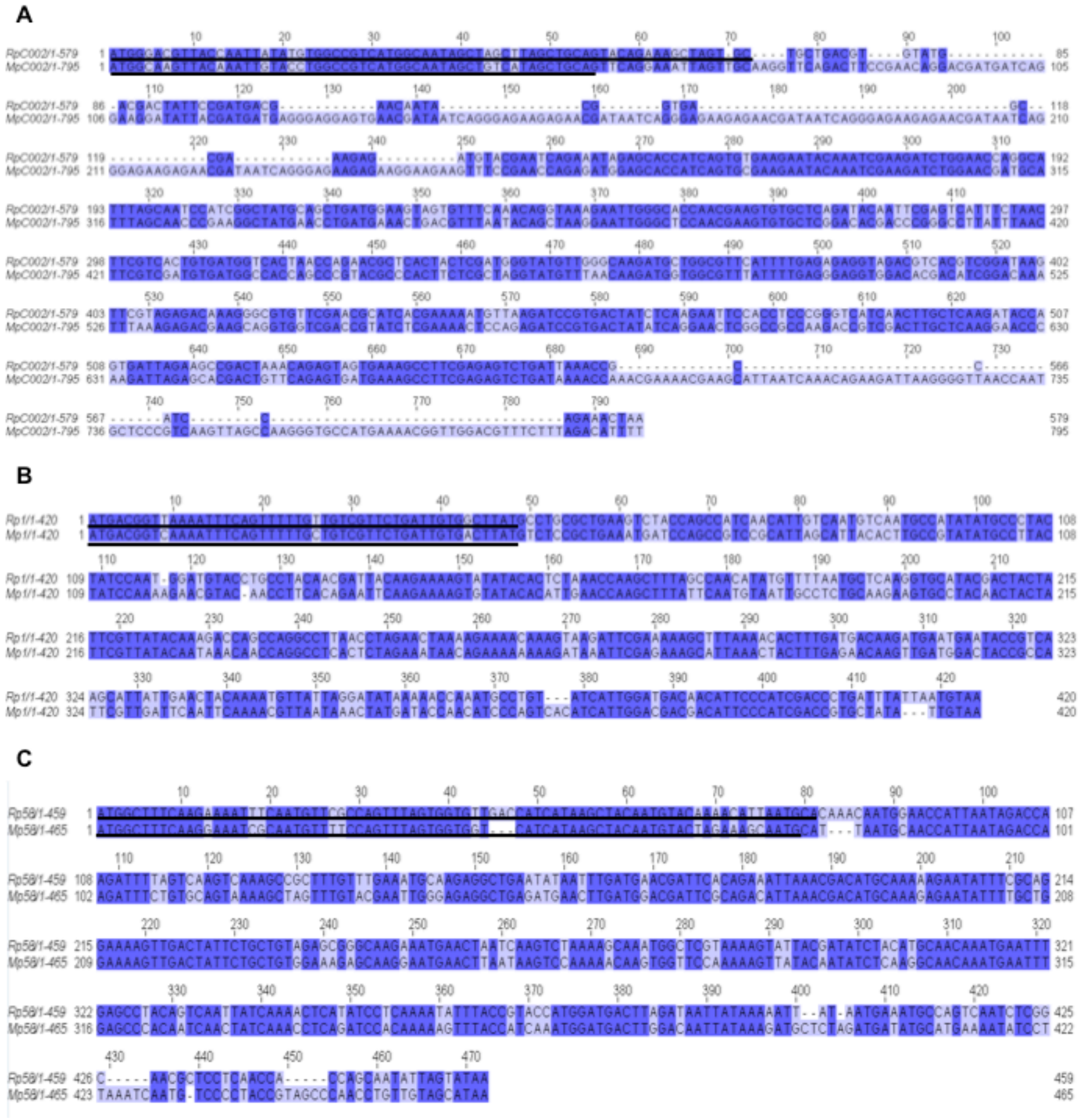

**Fig. S1. Pair-wise nucleotide sequence alignments of putative orthologous effectors from *Rhopalosiphum padi* and *Myzus persicae*.**

Alignments were generated using Jalview 2.10.4. The level of sequence conservation is indicated by dark (high conservation) to light purple colour (low conservation). Predicted signal peptide (SignalP 4.1) sequences are underlined in black.

**A**) RpC002/MpC002 alignment. **B**) Rp1/Mp1 alignment. **C**) Rp58/Mp58 alignment.

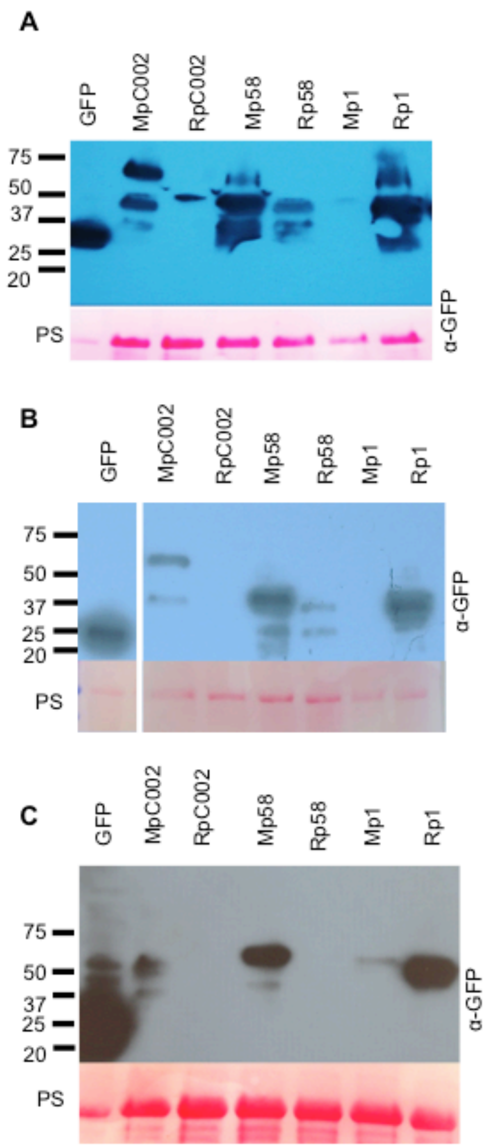


**Fig. S2. Western blots showing the expression of GFP and the GFP-effector fusion proteins in *Nicotiana benthamiana*.**

GPF and GFP-effector fusion proteins were transiently expressed in *N. benthamiana* by agroinfiltration. Leaf tissues were harvested 4 days post agroinfiltration for extraction and western blotting. Western blots showed the signal for GFP (±27kDa), GFP-MpC002 (±47kDa) GFP-RpC002 (±50kDa), GFP-Mp58 (±42kDa), GFP-Rp58 (±40kDa) and GFP-Mp1 (±47kDa), GFP-Rp1 (±40kDa). Lower panel shows Rubisco staining with Ponceau S (PS) showing equal amounts of protein loading except for the GFP sample (half volume loaded). The three different blots represent results from 3 biological replicates. For panel B the cropped lane showing the control GFP was from the same film as the GFP-fusion proteins shown on the right.

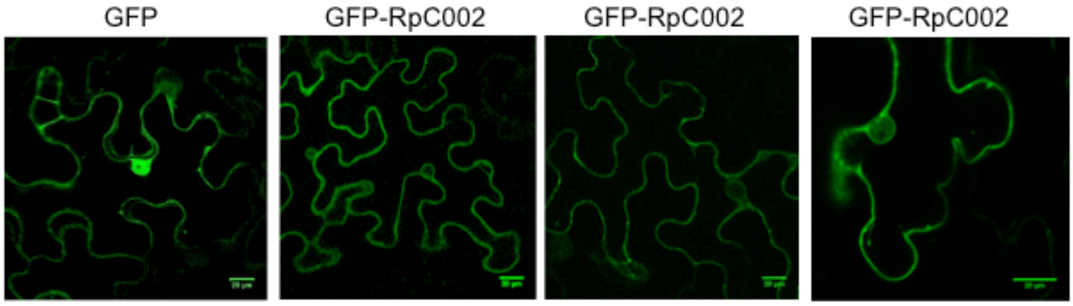


**Fig. S3. Localization of *Rhopalosiphum padi* effector RpC002 in *Nicotiana benthamiana*.** Additional confocal microscopy images showing free GPF and GFP-RpC002.

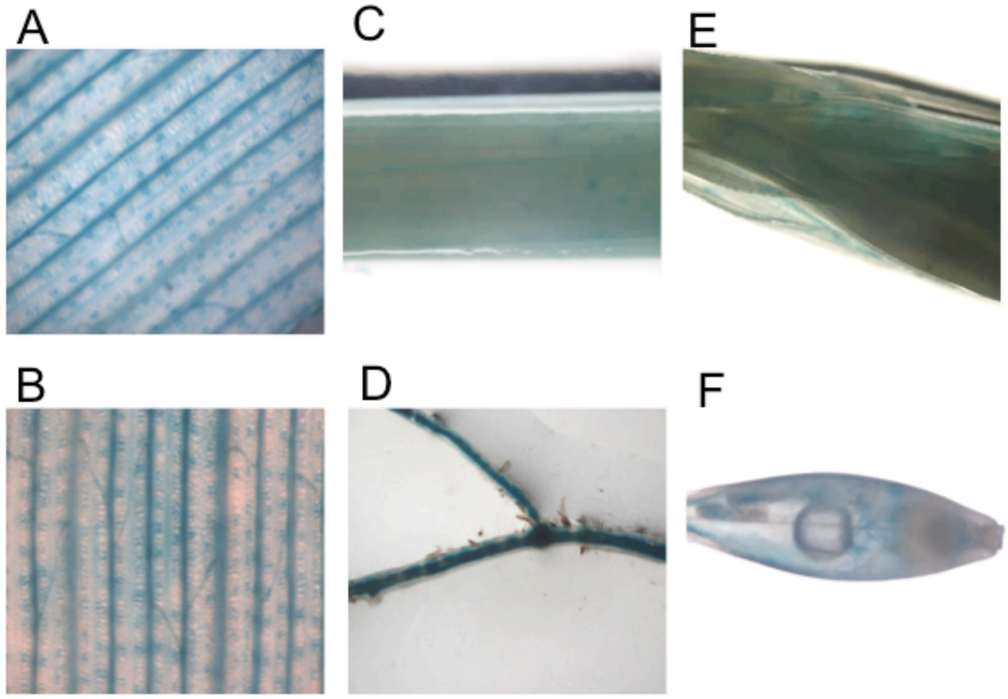

**Fig. S4. Expression of GUS (β-glucuronidase) under control of the maize ubiquitin promoter in different tissue of the barley transgenic line generated using pBRACT214:GUS.** GUS expression was visualized by staining barley tissue with X-gluc solution and overnight incubation at 37C. **A, D**) Leaf. **B**) Stem. **C**) Spike. **E**) Root. **F**) Grain.

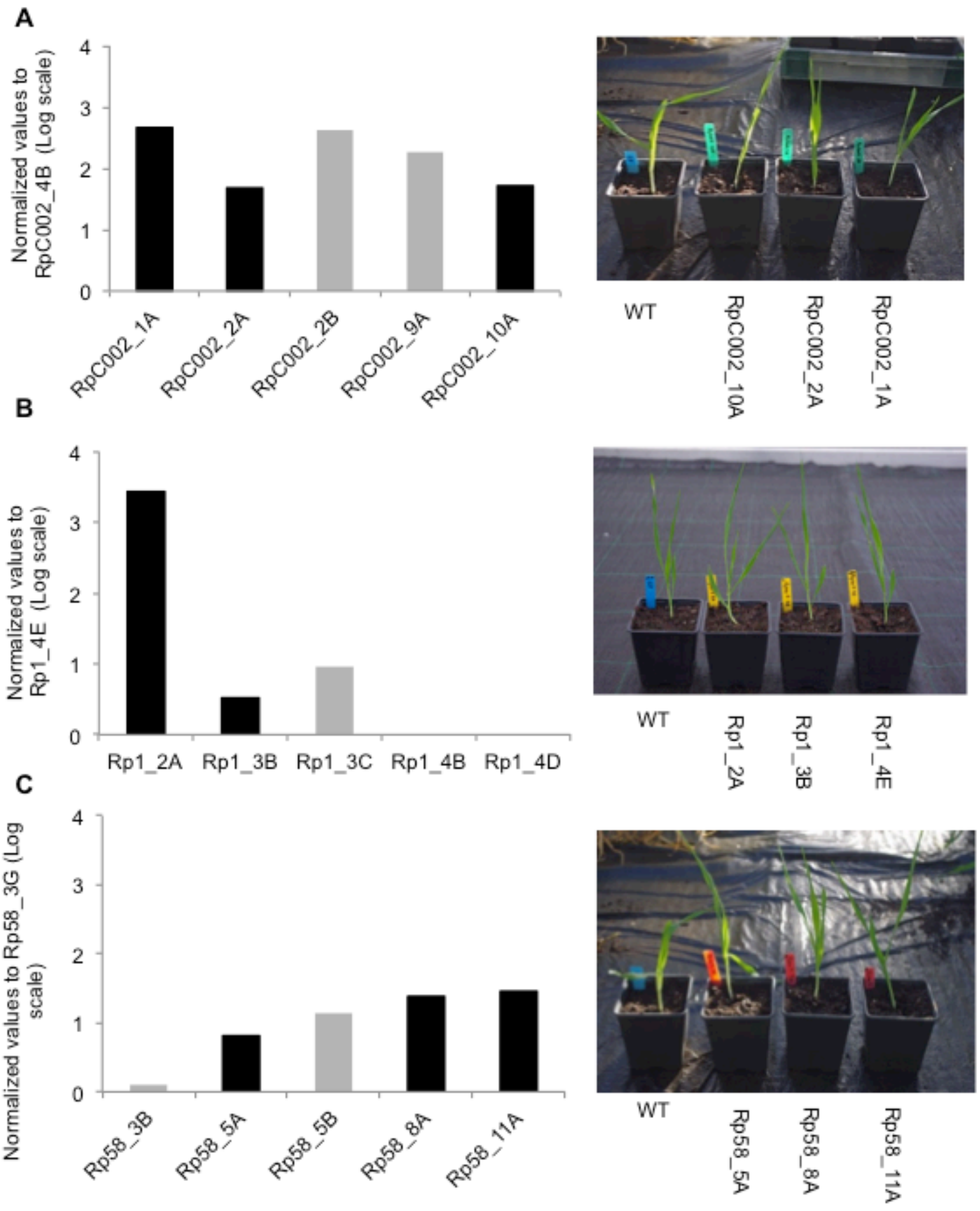


**Fig. S5. Effector transcript levels in transgenic barley lines expressing *Rhopalosiphum padi* effectors and plant phenotypes.** Expression was analysed by qRT-PCR on cDNA from selected transgenic barley lines in the first generation (T1). RNA was extracted from a single pool of 10-15 plants per genotype. Images on the right: phenotypes of homozygous 2 week-old transgenic lines selected for the aphid performance assays of each effector compared with the wild-type Golden Promise.

**A**) Normalized expression of RpC002 in transgenic lines relative to the line RpC002_4B on the left, and images of plants on the right.

**B**) Normalized expression of Rp1 in transgenic lines relative to the line Rp1_4E on the left and images of plants on the right.

**C**) Normalized expression of Rp58 in transgenic lines relative to the line Rp58_3G on the left and images of plants on the right.

In all graphs the lines used for the aphid performance assays (Fig. 5 and Fig. 6) are highlighted in black. Effector expression was not detected in the experimental controls, barley cv Golden Promise and the barley transgenic line transformed with the pBRACT214-GUS vector.

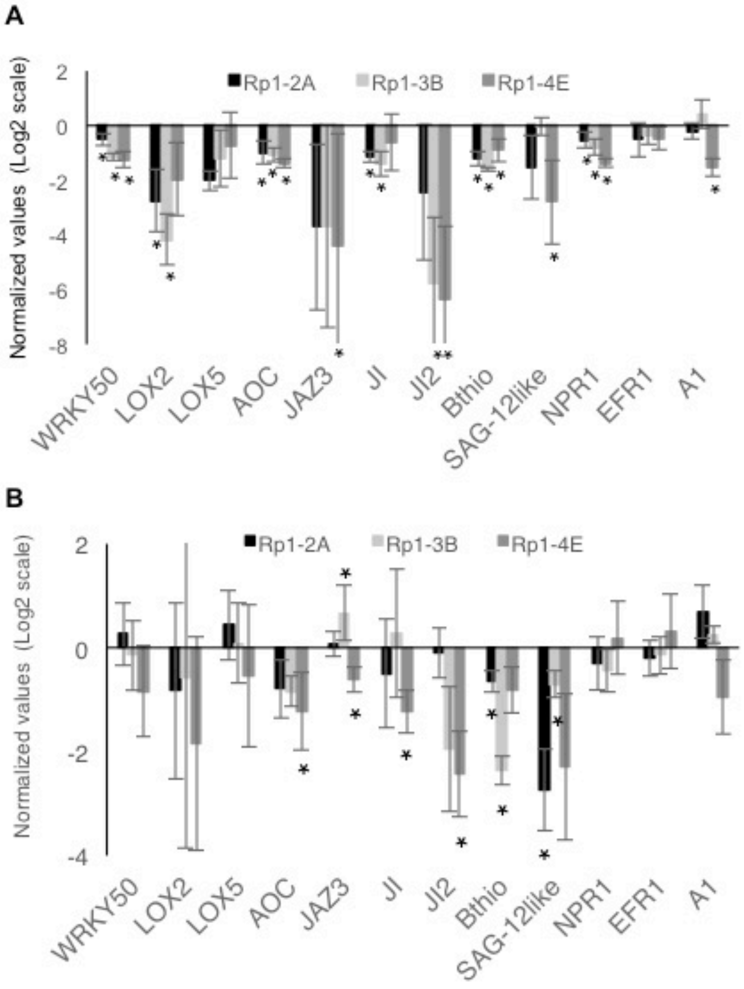


**Fig. S6. Defence-related gene expression in barley Rp1 lines after exposure to empty clip cages without *Rhopalosiphum padi*.**

Relative gene expression of defence-related/hormone-signalling genes was measured by qRT-PCR in control barley plants (cv. Golden Promise) and three independent barley lines expressing the *R. padi* effector Rp1.

**A)** Log-fold changes of barley gene expression upon 24h exposure to empty clip cages (no aphids) in the three independent Rp1 barley lines relative to control lines (WT=0).

**B)** Log-fold changes of barley gene expression upon 72h exposure to empty clip cages (no aphids) in the three independent Rp1 barley lines relative to control lines (WT=0).

All gene expression analyses were based on three independent biological replicates, and graphs represent mean expression. Genes are represented in the graphs are: *WRKY transcription factor 50* (*WRKY50*, MLOC_66204), *lipoxygenase 2* (*LOX2*, MLOC_AK357253), *lipoxygenase 5* (*LOX5*, MLOC_71948), *allene cyclase oxydase* (*AOC*, MLOC_68361), *jasmonate ZIM domain gene 3* (*JAZ3*, MLOC_9995), *jasmonate-induced gene* (*JI*, MLOC_15761), *jasmonate-induced gene 2* (*JI2*, MLOC_56924), *beta-thionin* (AK252675), *SAG12-like* (MLOC_74627.1), *non-expressor of pathogenesis-related 1-like* (*NPR1*, AM050559.1), the *ethylene-response factor 1* (*EFR1*, MLOC_38561) and *abscisic acid-inducible late embryogenesis abundant 1* (*A1*, MLOC_72442). Black bars represent gene expression levels in Rp1-2A lines, light grey bars represent gene expression levels in Rp1-3B lines and grey bars gene expression levels in Rp1-4E lines. Asterisks indicate significant differences between control plants (WT) and Rp1 transgenic lines (Wilcoxon Rank Sum Test, $p \leq 0.05$).

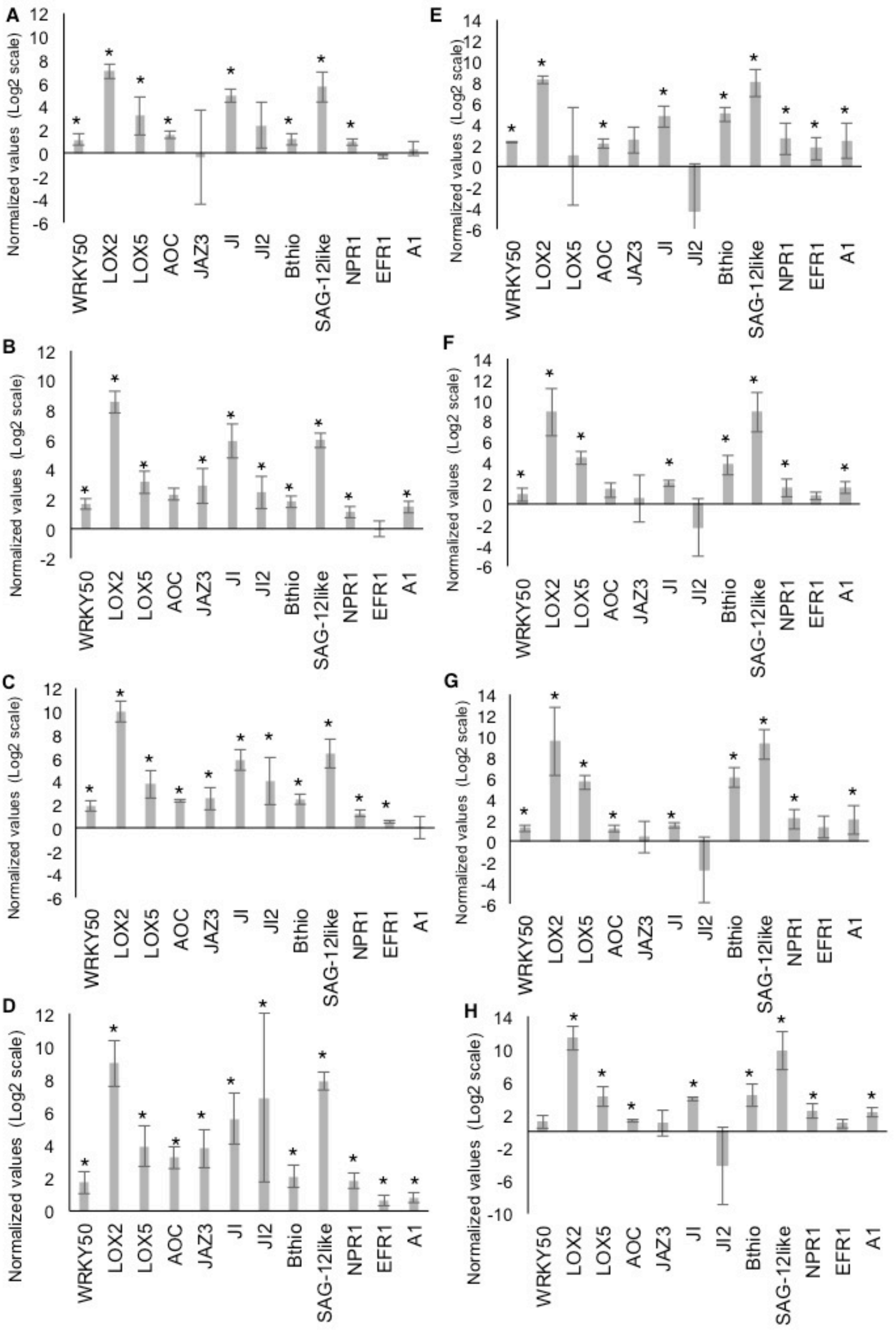


**Fig. S7. Defence-related gene expression in barley Rp1 lines after exposure to clip cages with *Rhopalosiphum padi*.**

Log-fold changes of defence gene expression measured by qRT-PCR in control barley plants (cv. Golden Promise) and three independent barley lines expressing the *R. padi* effector Rp1 upon infestation with *R. padi*.

**A)** Log-fold changes of defence gene expression upon aphid infestation of WT plants (clip cages with *R. padi* versus empty clip cages) after 24h.

**B)** Log-fold changes of defence gene expression upon aphid infestation of Rp1-2A plants (clip cages with *R. padi* versus empty clip cages) after 24h.

**C)** Log-fold changes of defence gene expression upon aphid infestation of Rp1-3B plants (clip cages with *R. padi* versus empty clip cages) after 24h.

**D)** Log-fold changes of defence gene expression upon aphid infestation of Rp1-4E plants (clip cages with *R. padi* versus empty clip cages) after 24h.

**E)** Log-fold changes of defence gene expression upon aphid infestation of WT plants (clip cages with *R. padi* versus empty clip cages) after 72h.

**F)** Log-fold changes of defence gene expression upon aphid infestation of Rp1-2A plants (clip cages with *R. padi* versus empty clip cages) after 72h.

**G)** Log-fold changes of defence gene expression upon aphid infestation of Rp1-3B plants (clip cages with *R. padi* versus empty clip cages) after 72h.

**H)** Log-fold changes of defence gene expression upon aphid infestation of Rp1-4E plants (clip cages with *R. padi* versus empty clip cages) after 72h.

All gene expression analyses were based on three independent biological replicates, and graphs represent mean expression. Genes are represented in the graphs are: *WRKY transcription factor 50* (*WRKY50*, MLOC_66204), *lipoxygenase 2* (*LOX2*, MLOC_AK357253), *lipoxygenase 5* (*LOX5*, MLOC_71948), *allene cyclase oxydase* (*AOC*, MLOC_68361), *jasmonate ZIM domain gene 3* (*JAZ3*, MLOC_9995), *jasmonate-induced gene* (*JI*, MLOC_15761), *jasmonate-induced gene 2* (*JI2*, MLOC_56924), *beta-thionin* (AK252675), *SAG12-like* (MLOC_74627.1), *non-expressor of pathogenesis-related 1-like* (*NPR1*, AM050559.1), the *ethylene-response factor 1* (*EFR1*, MLOC_38561) and *abscisic acid-inducible late embryogenesis abundant 1* (*A1*, MLOC_72442). Asterisks indicate significant differences between uninfested (empty clip cages) and infested (clip cages with aphids plants) (Wilcoxon Rank Sum Test, $p \leq 0.05$).

**Table S1. PCR primers used to clone the different effectors and qRT-PCR primers and probes used to quantify effector gene expression.**

| Primer name | Primer sequence | TaqMan Probe |
|---|---|---|
| AttL | **FP**_TCGCGTTAACGCCTAGCATGGATCTC/**RP**_GTAACATCAGAGATTTTGAGACAC | |
| AttL-TOPO | **FP**_CTCGCGTTAACGCTAGCATGGATGTT/**RP**_CCGTAACATCAGAGATTTTGAGACAC | |
| AttB | **FP**_GGGGACAAGTTTGTACAAAAAAGCAGGCT/**RP**_GGGGACCACTTTGTACAAGAAAGCTGGGT | |
| AttB-RpC002_F | GGGGACAAGTTTGTACAAAAAAGCAGGCTCC ATGGCTGACGTGTAT GAC GACTAT | |
| AttB-RpC002_R | GGGGACCACTTTGTACAAGAAAGCTGGGTCTTAGTTTCTGGATGGCGGTTTA | |
| AttB-Rp1 | (Rodriguez et al., 2017) | |
| TOPO-Rp58 | **FP**_CACCATGCAAACAATGGAACCATTAATAGAC/**RP**_TTATACTAATATTGCTGGTGGTTG | |
| Hv_Actin | **FP**_GCTTCAGATGCCCAGAGGT/ **RP**_GATGCCAGGAGCTTCCATAC | #11 |
| Hv_Pentatricopeptide | **FP**_CGGACCTCACCACCTTTAAC/ **RP**_AGGTCCCAGAACATGCACA | #16 |
| RpC002 | **FP**_CGCTCACTACTCGATGGGTAT/**RP**_CGTGACGTCTACCTCTCTCAAA | #159 |
| Rp1 | **FP**_TGCTCAAGGTGCATACGACT/**RP**_AGTGTTTTAAAGCTTTTTCGAATCTT | #20 |
| Rp58 | **FP**_CAGGAAAAGTTGACTATTCTGCTGTA/**RP**_GAGCCATTTGCTTTTAGACTTGA | #137 |
| Hv_Ubiquitin | **FP**_CATTCTCAATTCCCGAGCAG/**RP**_TTTTGGTGATGAAGCGGACT | |
| Hv_B_thionin | **FP**_TACTGGGTTTAGTTCTGGAGCAG/**RP**_ACGTGTCCTTGCAGCAACTT | |
| Hv_SAG12 like | **FP**_CAAGTGAGCAGTCGGACATC/**RP**_GGAAACGGTCCATCATCAGA | |
| Hv_NPR1 | **FP**_TTGATAACATCTAGAGGCAATGCT/**RP**_TGCGTGAAACTGTTCGAGAG | |
| Hv_EFR1 | **FP**_CTATATAATGATTGGGTGCATGTTG/**RP**_GGCATATGACCCAAGGTGTT | |
| Hv_A1 | **FP**_ATGGGAGGGGACAACACC/**RP**_GGAAATTAAGCGCGAACG | |
| Hv_LOX2 | **FP**_ATGTCCTATCCCACGACACC/**RP**_AGTGCGTCCTCAGCCAGT | |
| Hv_JAZ3 | **FP**_AGGAAAAGTGGTCGTGGTTG/**RP**_ATCTGGAGCAATCCGTTGAC | |
| Hv_WRKY50 | **FP**_TGTCCGTGCAAGGGGTAG/**RP**_ATTTTGGCGACGAGTATTCC | |
| Hv_AOC | **FP**_GCTACGAGGCCATCTACAGC/**RP**_AAGGGGAAGACGATCTGGTT | |
| Hv_JI | **FP**_GTGTACACCCCGTTGTTGC/**RP**_GCTCCTGCGACTGGTTATTC | |

| | |
|---|---|
| Hv_JI2 | **FP_**CAGCAGCGACTTCATTTACA/**RP_**ATGGTGTCGCAGACTATCCT |
| Hv_LOX5 | **FP_**AGGACTACCCGTACGCAGTG/**RP_**GGTGACCCATTTCTTGATGG |
